# Biosensor-enabled single-molecule enzyme activity profiling for function-based molecular counting

**DOI:** 10.64898/2026.09.18.752564

**Authors:** Toru Komatsu, Shosei Imai, Masaki Tajima, Yuki Sugiura, Takumi Iwasaka, Mayano Minoda, Yu Kagami, Kazufumi Honda, Takuya Terai, Reiko Tsuchiya, Ryosuke Kojima, Yasuteru Urano, Hiroyuki Kusuhara, Tadahaya Mizuno, Robert E. Campbell

## Abstract

Single-molecule enzyme activity analysis enables proteoform-resolved quantification of catalytically active enzyme molecules but remains limited by the need for enzyme-specific fluorogenic substrates, particularly for enzymes involved in central metabolism. Here we establish a biosensor-based framework that integrates genetically encoded fluorescent metabolite biosensors with single-molecule enzyme activity profiling (SEAP), enabling native enzymatic reactions to be monitored without enzyme-specific fluorogenic substrates. By employing different biosensors, the framework was applied to distinct metabolic enzymes, including lactate dehydrogenase and pyruvate kinase. Biosensor-based SEAP enables absolute quantification of active enzyme molecules, transforming conventional activity measurements (U/mL) into molecular abundance (molecules/mL). Applying this approach to single-cell functional proteoform analysis revealed functional heterogeneity in lactate dehydrogenase activity during T-cell activation. In addition, analysis of circulating enzymes showed that much of the plasma lactate dehydrogenase pool could be quantitatively accounted for by physiological erythrocyte turnover. These results establish genetically encoded fluorescent protein-based metabolite biosensors as modular optical transducers for single-molecule functional enzyme analysis across cellular and circulating samples.

## Introduction

Single-molecule enzyme activity profiling (SEAP) enables direct functional interrogation of individual enzyme molecules, revealing molecular heterogeneity that is obscured in conventional ensemble measurements^1,2^. By compartmentalizing enzymes into femtoliter-scale reactors, catalytically active molecules can be detected and counted individually, enabling highly sensitive quantification of functionally distinctive proteoforms^2,3^. Beyond resolving proteoform heterogeneity, this approach provides a conceptual advance in quantitative biochemistry by converting conventional enzyme activity measurements (U/mL) into molecular abundance (molecules/mL). This transformation provides a quantitative framework for linking enzyme activity to the abundance and dynamics of functional enzyme populations.

Despite these advantages, the scope of single-molecule enzyme assays remains limited because most rely on synthetic fluorogenic substrate analogues that must be individually designed and synthesized for each target enzyme^4–9^. This challenge is particularly acute for enzymes that catalyze interconversion of small-molecule metabolites, whose native substrates and products are often difficult to convert into fluorogenic reporters while preserving their recognition and reactivity toward the target enzyme^10^. Consequently, many enzymes central to metabolism remain inaccessible to SEAP^2^.

Here, we establish an approach that integrates genetically encoded fluorescent biosensors with SEAP for functional molecular counting. Genetically encoded fluorescent biosensors have been widely used to monitor intracellular ions and enzymatic activities, and their repertoire has recently expanded to diverse metabolites and second messengers^11–13^. While various designs of genetically encoded fluorescent biosensors have been reported^14^, those based on allosteric modulation of a single fluorescent protein typically provide the largest intensiometric responses^14^. Although these biosensors have been developed primarily to visualize biochemical dynamics in living cells, we reasoned that they could serve as modular optical transducers for microdevice-based single-molecule enzyme assays. By coupling native enzymatic reactions to biosensors that detect their small-molecule metabolic products, this strategy expands single-molecule functional analysis beyond enzymes accessible through conventional fluorogenic substrate design.

### A fluorescent protein biosensor enables proteoform-resolved functional analysis of LDH activity

As a proof of concept, we first applied the green fluorescent biosensor iLACCO^15,16^, which selectively responds to L-lactate, to the SEAP platform. Of the three reported variants with different affinities, we selected the high-affinity variant iLACCO1.2 with an apparent dissociation constant (*K*_d_) of 16.9 µM^16^. Because lactate is the direct product of the lactate dehydrogenase (LDH) reaction, we reasoned that iLACCO could provide a direct optical readout of native LDH activity. Previous single-molecule assays for LDH relied on detecting NADH during the reverse reaction (lactate-to-pyruvate conversion), which is thermodynamically unfavorable and does not readily distinguish between the physiologically relevant LDHA and LDHB isoforms^17^.

To adapt iLACCO to the SEAP platform, we optimized the assay conditions, including surfactant composition (**Figure S1**), to enable stable encapsulation of the biosensor within femtoliter reactors while preserving its fluorescence response (**Figure 2a, 2b, S2**). We also confirmed that GreenPy1Highest^18^, a fluorescent protein biosensor for pyruvate, retained a concentration-dependent response under the same loading conditions.

**Figure 1.**
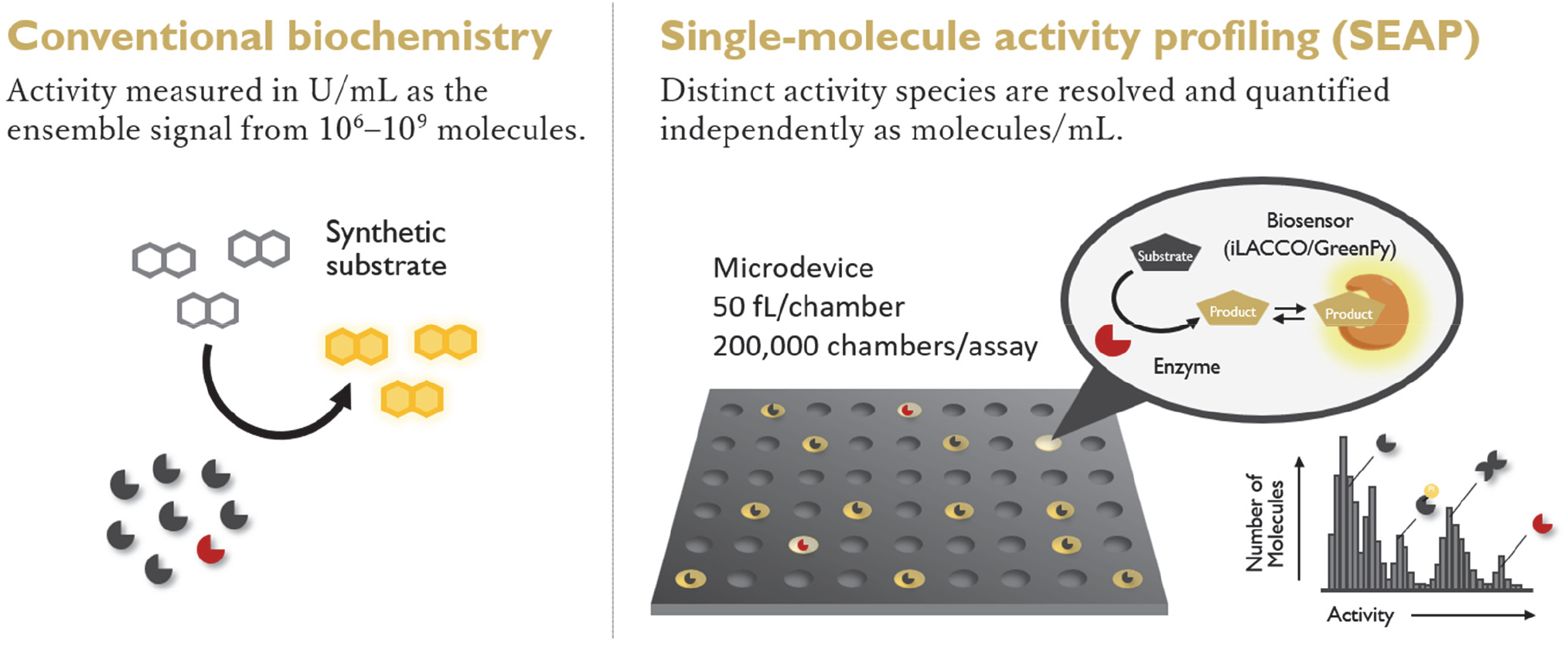
Schematic illustration of biosensor-enabled single-molecule enzyme activity profiling (SEAP). Comparison of conventional biochemical measurements of ensemble enzyme activity (left) and SEAP using femtoliter-scale microreactors (right). Enzymatic reaction products in individual reactors are detected using a genetically encoded fluorescent biosensor, and distinct activity species are resolved and quantified as molecules/mL.

**Figure 2.**
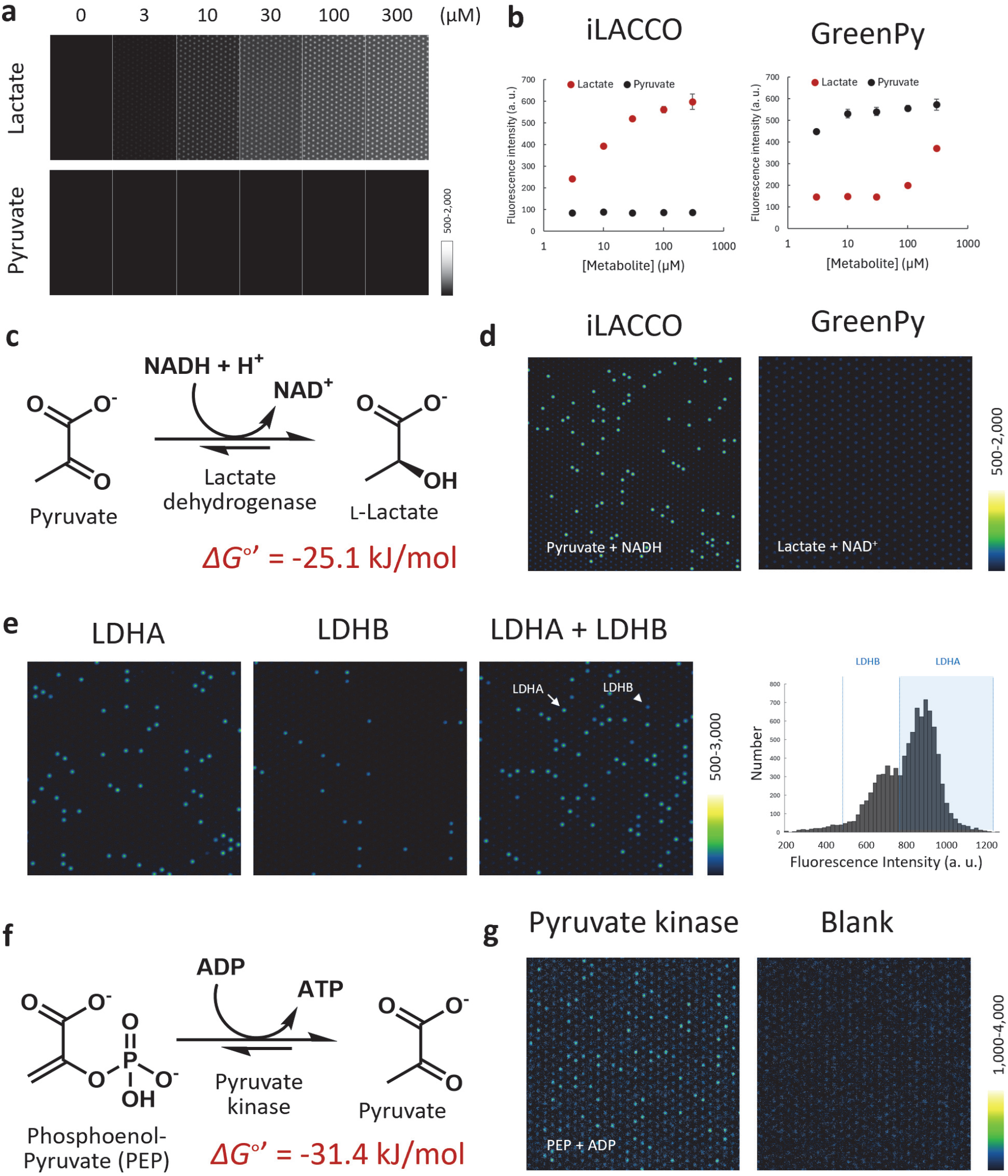
Construction of biosensor-based single-molecule enzyme activity assay. (a) Fluorescence images of microdevice loaded with iLACCO (10 μM) with varied concentrations of metabolites (L-lactic acid or pyruvic acid, 0-300 μM) in assay buffer (HEPES-Na buffer (100 mM, pH 7.4) containing CaCl_2_ (1 mM), MgCl_2_ (1 mM), DTT (100 μM), and Triton X-100 (150 μM)). (b) Quantification of fluorescence spots of microdevice loaded with iLACCO (10 μM) or GreenPy (10 μM) with varied concentrations of metabolites in assay buffer. (c) Reaction catalyzed by lactate dehydrogenase and its Gibbs free energy (*Δ*G°’) of the rightward reaction. (d) Fluorescence images of microdevice loaded with LDHB (0.1 ng/mL) with iLACCO (10 μM), pyruvate (500 μM) and NADH (500 μM) (left) or GreenPy (10 μM), L-lactate (30 μM) and NAD+ (100 μM) (right) in assay buffer and incubated at 25°C for 3 h. (e) Fluorescence images of microdevice loaded with LDHA (1 ng/mL) and/or LDHB (0.1 ng/mL) with iLACCO (10 μM), pyruvate (2 mM) and NADH (1 mM) in assay buffer and incubated at 25°C for 45 min. The histogram showing the intensity distribution of bright spots corresponding to LDHA and LDHB is shown. (f) Reaction catalyzed by pyruvate kinase and its Gibbs free energy (*Δ*G°’) of the rightward reaction. (g) Fluorescence images of microdevice loaded with pyruvate kinase (from rabbit muscle, 0.1 ng/mL) with GreenPy (10 μM), phosphoenol pyruvate (100 μM) and ADP (100 μM) in HEPES-Na buffer (100 mM, pH 7.4) containing KCl (100 mM), MgCl_2_ (10 mM), DTT (100 μM), and Triton X-100 (150 μM)) and incubated at 25°C for 2 h.

We then applied the optimized assay conditions to detect the activity of individual recombinant LDH molecules. Because LDHA and LDHB exhibit distinct kinetic properties to pyruvate^19^, we reasoned that their activities could be differentiated by tuning the pyruvate concentration. Indeed, increasing the pyruvate concentration progressively enhanced the difference between the fluorescence intensity distributions of recombinant LDHA and LDHB, enabling clear discrimination of the two isoforms at concentrations above 1 mM (**Figure 2e**). This distinction was also time-dependent: the fluorescence distributions of LDHA and LDHB became clearly separated within 30–60 min after reaction initiation, providing a temporal window for proteoform-resolved analysis (**Figure S3**). Positive reactors could be detected as early as 5 min after loading, whereas prolonged incubation eventually drove the signals toward a common plateau, consistent with the biosensor reporting the reaction equilibrium without enzymatically consuming the analyte (**Figure 1, Figure S3**). Together, these results established assay conditions that resolve individual LDHA and LDHB molecules based on their distinct catalytic properties. Together, these results established assay conditions that resolve individual LDHA and LDHB molecules based on their distinct catalytic properties. Under these optimized conditions, single-molecule counting also enabled highly sensitive detection of recombinant LDH, with a limit of detection (LOD) of 1.1 pg/mL (**Figure S4**). This sensitivity is substantially higher than that of conventional biochemical LDH assays, which typically have detection limits in the ng/mL range^4,20^, and is comparable to the sensitivity of conventional immunoassays for low-abundance circulating proteins^21^.

### Generality and design principles of protein biosensor-based SEAP

To examine the generality of biosensor-enabled SEAP, we next characterized GreenPy1Highest, a pyruvate-responsive fluorescent protein biosensor with a reported apparent *K*_d_ of 35 µM^18^. Under our assay conditions, GreenPy exhibited an apparent *K*_d_of approximately 10 µM in solution. Addition of components required for the enzymatic assay caused only modest changes in its apparent affinity, whereas confinement within the microdevice resulted in a more pronounced shift in its concentration-response profile (**Figure 2b, S2, S5**). Consistent with the previous report^18^, GreenPy also showed a detectable response to high concentrations of lactate (**Figure 2b, S5**). Because the concentrations required to elicit responses to pyruvate and lactate were substantially different, however, we reasoned that this selectivity could be accommodated by properly designing the assay conditions. We first replaced iLACCO with GreenPy to monitor the reverse reaction catalyzed by LDH. In this, the substrate concentration was limited to below 100 μM to minimize background fluorescence from lactate binding to GreenPy. However, in this condition, pyruvate accumulation was insufficient for robust single-molecule detection because the reaction equilibrium strongly favored lactate formation (**Figure 2d**). This result highlights an important design principle for biosensor-enabled SEAP: the enzymatic reaction must generate sufficient amounts of the biosensor-detectable product to reach the sensor’s response range within the confined reactor.

We therefore applied GreenPy to another enzyme, pyruvate kinase (PK), whose reaction directly generates pyruvate from phosphoenolpyruvate (PEP) and ADP (**Figure 2f**). PK is a central metabolic enzyme with important roles in cellular metabolism and cancer biology^22,23^. Under the concentration where the substrate PEP did not increase the background fluorescence (**Figure S5**), GreenPy enabled the detection of individual PK molecules in the microdevice (**Figure 2g**), demonstrating that a distinct enzyme reaction can be coupled to SEAP by exchanging the fluorescent protein biosensor.

### Single-cell functional profiling of LDH

Having established the analytical performance of the platform, we next explored whether its sensitivity and quantitative accuracy were sufficient for functional enzyme profiling at the single-cell level (**Figure 3a**). LDH is a central regulator of cellular metabolism and undergoes dynamic remodeling during immune cell activation^24,25^. Although single-cell transcriptomics have revealed extensive heterogeneity in LDHA expression during T-cell activation^24^, functional LDH activity is ultimately determined by the assembly of five homo- and heterotetrameric isoenzymes composed of LDHA and LDHB subunits, together with post-translational regulation. Direct measurement of catalytically active LDH molecules may therefore provide functional information that is not captured by transcript abundance alone.

**Figure 3.**
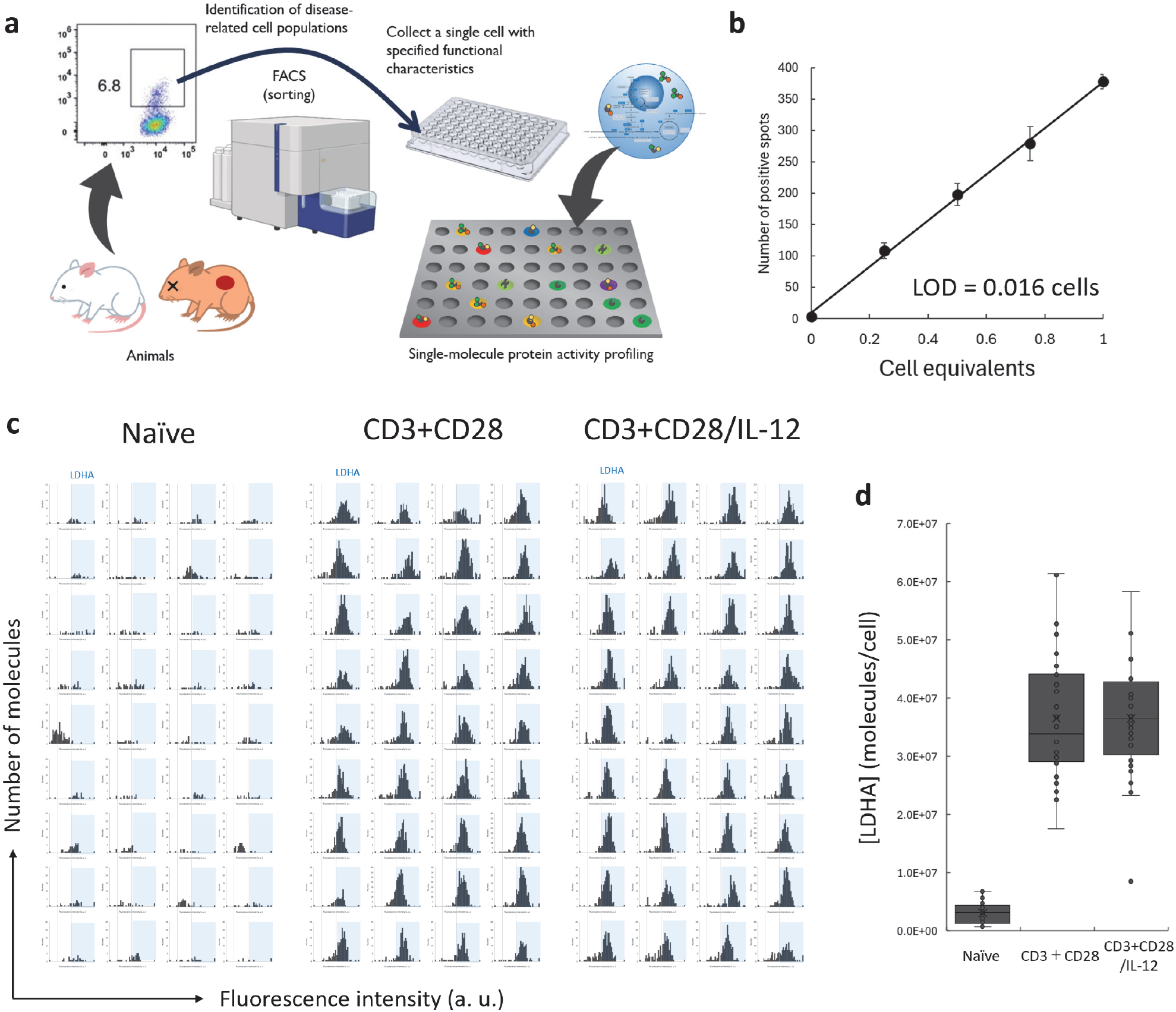
Construction of single-molecule LDH proteoform activity analysis. (a) Concept of single-molecule enzyme activity profiling of a single-cell. (b) Confirmation of the linearity of LDH spot number against cellular concentration. CD3/CD28-stilumated cells were used, and the data were acquired using cell lysates diluted to the indicated cell equivalents. n = 8 for 0 cell equivalents, and n = 4 for 0.2-1.0 cell equivalents. Error bars represent S. D. The limit of detection (LOD) was calculated as the cell equivalent corresponding to the mean signal of the blank plus three standard deviations (3 S. D.). (c) Histograms of LDH activity profiles of single cells. Each histogram corresponds to individual cells. Areas indicated by blue shading indicate activity populations corresponding to LDHA-major species. (d) Quantification of LDHA populations of cells in **Figure 3c**.

To evaluate the applicability of SEAP to single-cell functional proteoform analysis, we first determined its analytical sensitivity using lysates from activated T cells. The assay exhibited a LOD of 0.016 cell equivalents while maintaining excellent linearity well below the single-cell level (**Figure 3b**). This level of quantitative performance is notable because accurate protein quantification from subcellular inputs remains a major challenge for current single-cell omics approaches^26–28^.

We then profiled LDH activity in individual naïve e T cells and T cells subjected to anti-CD3/CD28 stimulation in the absence or presence of IL-12, conditions known to induce distinct functional states of CD8^+^ T cells^29–31^. Following activation, cells were isolated individually by flow cytometry (**Figure 3a**), and SEAP revealed substantial cell-to-cell heterogeneity in the abundance of active LDH molecules across all populations. These populations primarily expressed LDHA-dominant LDH species^25^, and mean active LDH abundances were 3.2 × 10^6^, 3.4 × 10^7^, and 3.7 × 10^7^ molecules per cell in naïve e, CD3/CD28-stimulated, and CD3/CD28/IL-12-stimulated cells, respectively (**Figure 3c, 3d**). It has been reported that the elevation of LDHA transcripts upon T cell activation is less than five-fold^24^, whereas our functional measurements revealed a greater than tenfold increase in active LDH molecules. The larger change observed at the functional protein level may reflect additional layers of regulation beyond transcription, including translation, protein stability, tetramer assembly, and post-translational regulation.

### Quantitative analysis of circulating LDH reveals physiological erythrocyte turnover

We next applied the platform to analyze circulating LDH species. Consistent with previous biochemical studies^32^, LDHB predominated in human plasma. Whereas conventional measurements report total LDH activity in U/mL, SEAP directly quantified the abundance of catalytically active LDH molecules, revealing a median plasma concentration of 1.65 × 10^12^ molecules/mL across 95 healthy individuals (**Figure S6**). This corresponds to a nanomolar concentration of catalytically active LDH in healthy circulation, which appeared unexpectedly high compared with that of many circulating proteins with established mechanisms of secretion into the bloodstream^21,33^. LDH is an established biomarker of tissue injury, including myocardial injury^32,34,35^, but the source of basal circulating LDH in healthy individuals remains unclear. Because LDH lacks an established pathway for active secretion into the circulation^36–38^, we hypothesized that circulating LDH may reflect constitutive cell turnover. We considered erythrocytes, red blood cells (RBCs), as a major potential source because they are the most abundant cell population undergoing continuous turnover^39^, accounting for approximately 65% of total daily cell death, corresponding to approximately 2.1 × 10^11^ cells dying per day. Although most erythrocytes undergo cell death by eryptosis and are removed by splenic macrophages, it is considered that 10–20% are lost intravascularly, releasing intracellular proteins into the circulation^40,41^. We therefore constructed a quantitative mass-balance model to ask whether circulating LDH could be explained by the release of intracellular LDH from erythrocytes undergoing intravascular cell death (**Figure 4a**).

**Figure 4.**
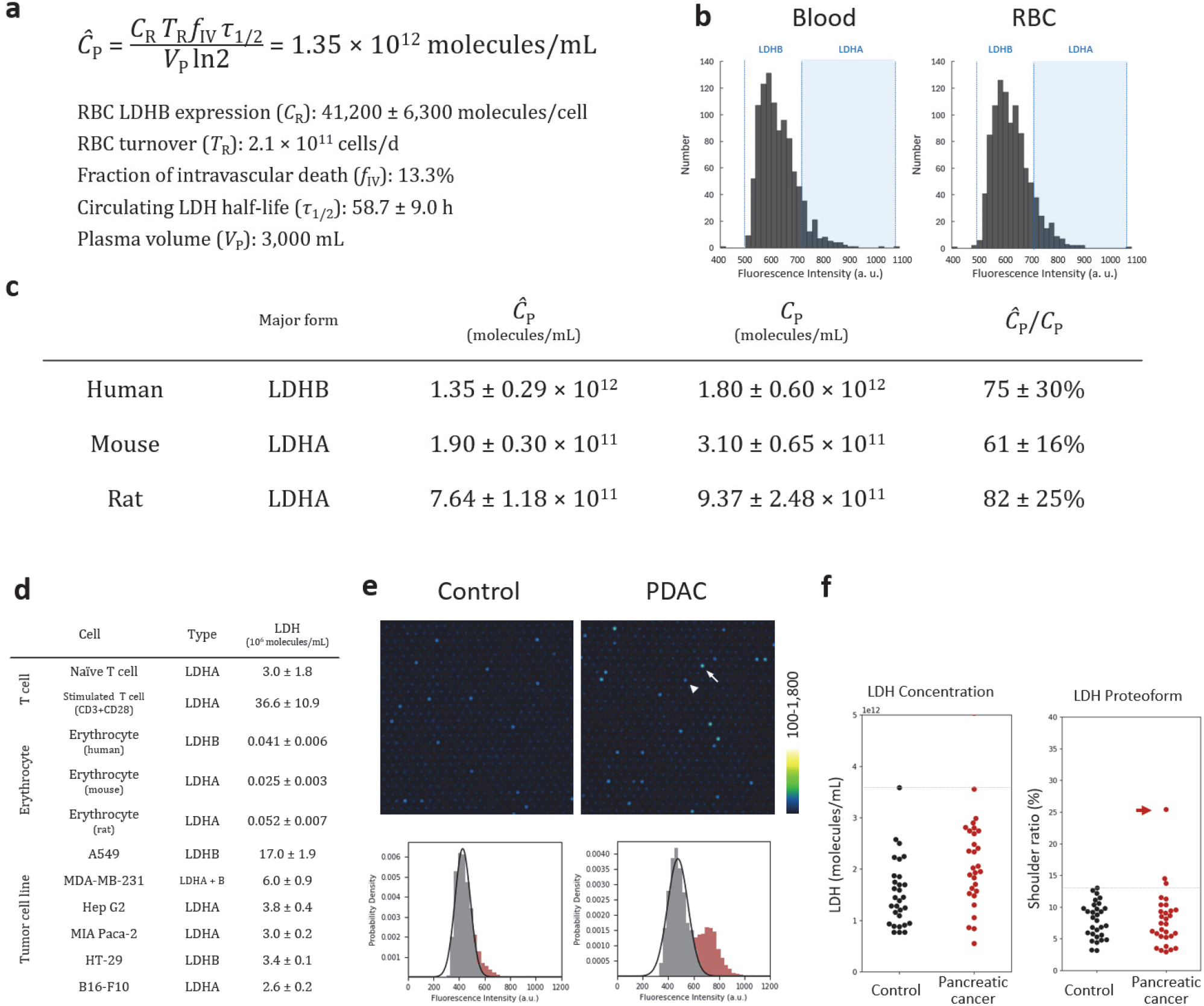
Estimation of LDH concentrations generated from intravascular erythrocyte death. (a) Equation to calculate the estimated plasma concentration of LDH released from intravascular cell death (*Ĉ*_P_). RBC LDHB expression (*C*_R_) was based on the measurement, and RBC turnover (*T*_R_)^39^, fraction of intravascular death (*f*IV)^40^. plasma half-lives of circulating LDH (*τ*_1/2_)^42^, and plasma volume (*V*_P_)^40^ are from the literature. (b) Activity histograms of human plasma (left) and erythrocyte (right). The measurement was performed by mixing plasma samples (1/2,500 dilution) or RBC lysate (1/10,000 dilution), iLACCO (10 μM), pyruvate (2 mM) and NADH (1 mM) in assay buffer, loading into microdevice, and incubated at 25°C for 45 min. (c) Comparison of calculated *Ĉ*_P_ and LDH concentration (*C*_P_) measured using plasma samples of human, mouse, and rat. (d) Concentration of LDH (molecules/cell) and major isoforms in various cell types. The activity histograms of model cancer cells are shown in **Figure S10**. (e) Representative example of altered plasma LDH activity profiles observed in PDAC patients. In fluorescence images, a white arrow indicates LDHA and a white arrowhead indicates LDHB. Activity species exceeding the single Gaussian distribution of LDHA (marked as red) is assigned as shoulder peak, and the number of molecules assigned as shoulder peak per total LDH is assigned as shoulder ratio (%). Histograms of whole samples are shown in **Figure S9** and **S10**. (f) Results of total LDH number (left) or shoulder ratio (%) of 1^st^ cohort. n = 30 for healthy subjects and n = 30 for pancreatic cancer patients. The result of 2^nd^ cohort (n = 95 for healthy subjects and n = 38 for pancreatic cancer patients) is shown in **Figure S11** and **S12**. A red allow indicates the patient picked up in (e).

The LDH activity profile of purified human erythrocytes closely recapitulated that of plasma (**Figure 4b**), and each erythrocyte contained approximately 41,000 active LDH molecules. Using this experimentally determined molecular abundance together with reported erythrocyte turnover rates^39^ and plasma LDH half-lives^42,43^, we estimated the steady-state plasma LDH concentration expected from erythrocyte turnover (**Figure S7**). Assuming that 10–20% of erythrocytes undergo intravascular cell death, the predicted steady-state concentration (*Ĉ*_P_) was 1.02-2.04 × 10^12^ molecules/mL closely encompassing the measured median concentration (*C*_P_) of 1.65 × 10^12^ molecules/mL (**Figure 4c, S6**). Thus, the data suggested that physiological erythrocyte turnover could account for a substantial fraction of circulating LDH under healthy conditions.

We next tested this model across species. In contrast to the LDHB-dominant profiles observed in humans, LDHA predominated in erythrocytes of both mice and rats, and their activity profiles closely matched those observed in the corresponding plasma samples (**Figure S8**). Accounting for species- and isoform-specific plasma half-lives, the predicted-to-measured concentration ratios (*Ĉ*_P_/*C*_P_) were 75±30%, 61±16%, and 82±25% in human, mice and rats, respectively (**Figure 4c**). We further examined whether viable cells released active LDH independently of cell death. Across erythrocytes from all three species and six cancer-derived cell lines (lung, breast, liver, pancreas, and colorectal cancer cells and melanoma cells), extracellular release of active LDH was negligible (less than 1% of cellular LDH detected in 1 h), supporting that the release of LDH into circulation is unlikely to occur without cell death (**Figure 4d, S9**). Together, these results support constitutive erythrocyte turnover as a major source of basal circulating LDH.

### Disease-associated LDH profiles enable quantitative inference of lclular origin

We next asked whether disease-associated alterations in circulating LDH profiles could be interpreted using the same quantitative framework. Plasma from patients with pancreatic cancer (PC) identified a patient with a distinct circulating LDH activity profile characterized by an LDHA-associated shoulder (**Figure 4e, S10, S11**). This distinct phenotype was first identified in one patient in a cohort of 30 PC and 30 control samples (**Figure 4e, 4f, S10**) and was subsequently observed again in a larger independent cohort of 95 PC and 38 control samples (**Figure S11, S12a**), supporting that it represents a reproducible PC-associated blood phenotype rather than an interinstitutional variation.

Three plausible sources of circulating LDHA—pancreatic cancer cells, activated T cells, and hepatocytes—were considered because the corresponding model cells mainly expressed LDHA (**Figure 4d, S8**), but quantitative mass-balance analysis argued against the first two sources. The patient with the most prominent LDHA increase had approximately 5.6 × 10^11^ molecules/mL of LDHA in plasma. Given the short plasma half-life of human LDHA (approximately 10 h^44^), maintaining this concentration would require the continuous release of approximately 2.8 × 10^15^ LDHA molecules per day, equivalent to lysis of 9.3 × 108 MIA PaCa-2-equivalent pancreatic cancer cells per day (**Figure 4d**), or approximately 1 g of tumor tissue daily. This magnitude of cell loss makes the tumor itself an unlikely dominant source. Activated T cells contained more LDHA than pancreatic cancer cells (**Figure 4d**), but T cells comprise less than 10% of tumor cellularity in pancreatic cancer^45^, also making T-cell death quantitatively insufficient to explain the observed signal. In contrast, the corresponding estimate for hepatocytes was 7.3 × 108 cells per day, based on LDHA abundance in Hep G2 cells (**Figure 4d**). This represents approximately 0.3% of the estimated 2.4 × 10^11^ cells in the human liver and falls within the reported range of physiological hepatocyte turnover (< 0.4% per day)^39^. Biliary obstruction, which occurs in a subset of patients with pancreatic cancer^46^, can additionally induce localized hepatocyte injury and necrosis^47^. Consistent with this possibility, circulating GGT1 activity, a clinical marker associated with cholestasis and liver injury^48^, was elevated in the patient with the prominent LDHA increase (**Figure S12b**). Although the hepatocyte origin of circulating LDHA cannot be definitively established from the current data, they illustrate how absolute molecular counting constrains the cellular origins that are quantitatively compatible with a circulating enzyme signal.

## Discussion

In this study, we established a framework for coupling genetically encoded fluorescent biosensors to microdevice-based single-molecule enzyme activity profiling (SEAP) for quantitative analysis of active enzyme proteoforms. Unlike conventional single-molecule enzyme assays, which typically rely on enzyme-specific fluorogenic substrates^1,8^, this strategy enables single-molecule analysis of native enzymatic reactions through biosensor-based detection of their products. Given the rapidly expanding repertoire of genetically encoded biosensors for metabolites and second messengers^11–13^, this framework has the potential to substantially broaden the range of enzymes accessible to single-molecule functional analysis.

A second conceptual advance of biosensor-enabled SEAP is the direct quantification of catalytically active enzyme molecules. Clinical enzymology has traditionally quantified enzymes by catalytic activity, expressed as units per mL (U/mL). While such measurements have proven invaluable in both research and clinical practice, activity units conflate molecular abundance with the catalytic properties of the underlying enzyme population and therefore do not directly report how many functional enzyme molecules are present. By detecting individual active enzymes, SEAP resolves these two quantities, converting ensemble activity into the absolute abundance of catalytically active molecules (molecules/mL). This distinction becomes particularly important when enzyme populations comprise functionally distinct proteoforms or when molecular abundance must be related quantitatively to the cellular processes that generate them. Single-cell analysis illustrates a complementary consequence of functional molecular counting. T-cell activation produced a greater than tenfold increase in active LDH molecules per cell, exceeding the change previously observed at the transcript level^24^. Although the molecular mechanisms underlying this discordance remain to be established, it underscores that transcript abundance alone does not specify the abundance of catalytically active enzyme molecules. Measurements at the functional protein level can integrate multiple layers of regulation, including protein abundance, oligomeric assembly and post-translational regulation, providing information complementary to single-cell transcriptomic measurements^49^.

The application to circulating LDH illustrates how absolute functional molecular counting can connect a circulating enzyme signal to its underlying cellular source. LDH has long been regarded as a classical biomarker of tissue injury, yet the basis for its relatively high abundance in healthy circulation has remained poorly understood^32,34,35^. By combining experimentally measured LDH molecules per erythrocyte with reported cellular turnover and plasma clearance kinetics, we found that physiological erythrocyte turnover could account for a substantial fraction of circulating LDH. The agreement between predicted and measured concentrations across humans, mice and rats, despite species-specific differences in the predominant LDH isoform, supports this quantitative interpretation. Although the model relies on reported turnover rates and estimated plasma half-lives, it illustrates how molecular counting can transform a circulating enzyme concentration into a testable mass-balance model of its production and clearance. Such liquid-derived information on functional enzyme dynamics (LiFE) should become increasingly precise with the incorporation of additional cell types, subtype-specific kinetics and proteoform-dependent clearance rates.

Several limitations and design considerations should be acknowledged. First, successful implementation of biosensor-enabled SEAP requires matching the catalytic properties and thermodynamics of the target reaction to the affinity and dynamic range of the biosensor and the volume of the microreactor. As illustrated by the unsuccessful coupling of GreenPy to LDH, sufficient product must accumulate within the reactor to generate a detectable biosensor response. This requirement may become particularly important for slow-turnover enzymes, which will require sensors responsive at lower analyte concentrations^2^. Whereas biosensors for live-cell imaging are typically optimized for physiological intracellular concentrations^13,15^, single-molecule enzymology may therefore motivate a complementary direction in biosensor engineering towards high affinity for sensitive detection of low concentrations. Finally, the physiological turnover model presented here represents an initial approximation. Direct measurements of proteoform-specific plasma half-lives, release mechanisms and cell type-specific turnover rates will be necessary to establish more complete models of circulating enzyme homeostasis. Likewise, quantitative modeling of disease-associated LDH profiles can constrain their potential cellular origins, but these assignments will require independent experimental validation. Nevertheless, the ability to express circulating enzyme abundance in molecular units provides a quantitative basis on which such mechanistic models can be constructed and tested.

## Conclusion

In summary, biosensor-enabled SEAP connects two established technological frameworks—genetically encoded fluorescent biosensors and single-molecule enzymology—to enable functional molecular counting of native enzyme reactions. By moving from ensemble activity units to absolute numbers of active enzyme molecules, the approach enables functional proteoform analysis from individual cells and quantitative interpretation of circulating enzyme abundance in terms of cellular turnover. As the repertoire and performance of genetically encoded biosensors continue to expand, this framework should broaden the range of biochemical reactions accessible to single-molecule functional analysis and provide new opportunities to connect molecular enzyme activity with cellular and physiological processes.

## Methods

### Preparation of plasma samples from human subjects and ethics statement

For the studies conducted in The University of Tokyo (1^st^ cohort), plasma samples were obtained from the Platform for Evaluating Biomarkers of Cancer Early Detection (P-EBED), with the Program for Promotion of Fundamental Studies in Health Sciences conducted by the National Institute of Biomedical Innovation of Japan, Health and Labour Sciences Research Grants from the Ministry of Health, Labor and Welfare of Japan, and P-CREATE of the Japan Agency for Medical Research and Development (AMED)^1–3^. Ethical approval for the study was obtained from the central ethics committees of Nippon Medical School (M-2021-002) and the ethical committee of Nippon Medical School (A-2020-032 and A-2020-044).

For the studies conducted in Cosomil, Inc. (2^nd^ cohort), plasma samples were collected from the Okayama University Hospital Biobank (Okadai Biobank) and Japan Institute for Health Security Biobank, Japan. For samples collected at Cosomil, Inc., blood was drawn into EDTA-containing tubes. The collected blood samples were generally centrifuged at room temperature at 2,200 × *g* for 10 min to separate plasma. Ethical approval for the study was obtained from Kobe University Hospital (B230079), Craif Institutional Review Board (IEF-24012401), and PHRF-IRB (24B0001) (for experiments conducted in Cosomil, Inc.). Disease stage was assigned according to the Union for International Cancer Control (UICC) TNM classification, 8^th^ edition.

### Preparation of red blood cells from human subjects and ethics statement

Experiments were performed in Cosomil, Inc. Ethical approval for the study was obtained from Kobe University Hospital (B230079), Craif Institutional Review Board (IEF-24012401), and PHRF-IRB (24B0001) (for experiments conducted in Cosomil, Inc.). Blood was collected from the antecubital vein into EDTA-containing blood collection tubes. For erythrocyte isolation, 300 μL of whole blood was diluted twofold with saline (n = 2 preparations per individual), and the diluted blood was carefully layered onto 600 μL of the OptiPrep density-gradient medium (*d* = 1.095 g/mL) without disturbing the interface. Samples were centrifuged at 800 × *g* for 20 min at 25°C. The erythrocyte pellet was washed with saline and resuspended in 200 μL saline, followed by three additional washes with 1 mL saline (800 × *g*, 1 min). Erythrocyte was resuspended in 1 mL saline, and the cell counting was performed using cell counter (Countess 3 FL, Thermo). For preparation of erythrocyte lysates, an equal volume of 2× lysis buffer (10 mM HEPES, pH 7.4, 3 mM Triton X-100) was added to the erythrocyte suspension. After incubation for 10 min, samples were centrifuged at 15,000 × *g* for 1 min, and the supernatant was collected, aliquoted, and flash frozen using liquid N_2_ for subsequent analysis. To evaluate extracellular LDH release, erythrocyte suspensions were incubated at 37°C, and extracellular fractions were collected at 0 and 1 h by centrifugation at 5,000 × *g* for 1 min.

### Preparation of plasma samples from animals and ethics statement

Ethical approval for the study using animals was obtained from Animal Care and Use Committee of The University of Tokyo (P4-21, P31-9). Six-week-old male C57BL/6JJcl mice were purchased from CLEA Japan (Tokyo, Japan). After acclimatizing for more than five days, mice were euthanized and the blood was collected through an inferior vena cava into 1.5 mL tube containing 1.5 µL heparin (Yoshindo Inc, Japan). The collected blood sample was centrifuged (1,700 × *g*, 4°C for 15 min) for plasma separation, and the resulting plasma was stored at −80°C until analysis. Male SD rats (n = 6) were purchased from Japan SLC, Inc. (Shizuoka, Japan). At eight weeks of age, blood samples were collected from the tail vein without anesthesia into 1.5 mL tubes containing 1.5 μL of heparin sodium (10,000 units/10 mL, Heparin Sodium Injection “AY”, Yoshindo Inc., Japan). Approximately 200 μL of blood was collected from each rat without fasting. The collected blood samples were centrifuged (1,700 × *g*, 4 °C for 15 min) for plasma separation, and the resulting plasma was stored at 313:14280°C until analysis.

### Harboe method

Plasma samples (1.5 μL) was loaded onto Nanodrop One (Thermo), and absorbance at 380, 415, and 450 nm were measured. Free hemoglobin concentration (fHb) was calculated as fHb = 83.6 × (2 × A_415_ 313:142A_380_ 313:142A_450_) (mg/dL). Hemolysis % was calculated from total hemoglobin concentration = 14 g/dL for mice and 15 g/dL for rat^4^. Samples with hemolysis% > 1% were not included in the analysis.

### Preparation of red blood cells from animals and ethics statement

Ethical approval for the study using animals was obtained from Animal Care and Use Committee of The University of Tokyo (P4-21, P31-9). Six-week-old male C57BL/6JJcl mice and eight-week-old male Wistar rats were purchased from CLEA Japan (Tokyo, Japan). Blood was collected by cardiac puncture under anesthesia into heparin-coated collection tubes. For erythrocyte isolation, 300 μL of whole blood was diluted twofold with saline, and the diluted blood was carefully layered onto 600 μL of OptiPrep density-gradient medium (*d* = 1.090 g/mL) without disturbing the interface. Samples were centrifuged at 800 × *g* for 20 min at 25°C using a soft brake. The erythrocyte pellet was washed with saline and resuspended in 200 μL saline, followed by three additional washes with 1 mL saline (800 × *g*, 1 min). Erythrocytes were then resuspended in 1 mL saline, and cell counting was performed using a cell counter (Countess 3 FL, Thermo Fisher Scientific). For preparation of erythrocyte lysates, an equal volume of 2× lysis buffer (10 mM HEPES, pH 7.4, 3 mM Triton X-100) was added to the erythrocyte suspension. After incubation for 10 min, samples were centrifuged at 15,000 × *g* for 1 min, and the supernatant was collected, aliquoted, and flash-frozen in liquid N2 for subsequent analysis. To evaluate extracellular LDH release, erythrocyte suspensions were incubated at 37°C, and extracellular fractions were collected at 0 and 1 h by centrifugation at 5,000 × *g* for 1 min.

### Enzymes

Lactate dehydrogenase A from *Homo sapiens* was purchased from R&D systems (9158-HA, Lot #DFYD0223051). Lactate dehydrogenase B from *Homo sapiens* was purchased from R&D systems (9205-HB, Lot #DGGN0123011). Pyruvate kinase from rabbit muscle was purchased from Sigma-Aldrich (P7768, Lot #0000526782).

### Preparation of protein biosensors

The genes encoding iLACCO1.2 and GreenPy1Highest with a poly-histidine tag on the N-terminus were expressed from the pBAD vector in *E. coli* strain DH10B (Thermo Fisher Scientific). The bacteria were cultured in LB media with 100 μg mL^−1^ ampicillin at 37°C until the OD600 reached 0.6. Expression was then induced by adding 0.02% L-arabinose and the cells were cultured overnight at room temperature. Cell pellets were lysed with a cell disruptor (Branson), and proteins were purified by Ni-NTA affinity chromatography (Wako). Following purification, the buffer was exchanged by Amicon Ultra-15 30kDa (Millipore) to 30 mM MOPS buffer (pH 7.2) with 100 mM KCl.

### Purification of protein biosensors using gel filtration chromatography

For analysis of single-cell LDH, protein biosensor was purified by size-exclusion chromatography using a Superdex 75 Increase 10/300 GL column on an ÄKTA Explorer system (Cytiva). The column was equilibrated and operated with 30 mM MOPS and 100 mM KCl (pH 7.2) as the running buffer.

### Fluorescence microscopy

Fluorescence images were acquired by fluorescence microscope (Ti2, Nikon) equipped with a 20× dry objective lens (S Plan Fluor LWD 20×C; NA = 0.70, WD = 2300 μm) and a motorized stage. The assay was performed using a solution containing dsSiR^5^ (1 μM) as the internal standard, and the focus was adjusted using its fluorescence. Images were acquired in tile scan mode of 3×4 (with overlap of 1%) with perfect focus. For epifluorescence microscopy, fluorescence images were acquired using sCMOS camera (ORCA-Fusion C14440, Hamamatsu Photonics) under excitation with white LED illumination unit (X-Cite Xylis, Opto Science). The excitation and emission filters used were FITC (for GFP-based sensors, mirror = 505 nm, Ex. = 465-495 nm, Em. = 512-558 nm) and mCherry (for dsSiR, mirror = 600 nm, ex. = 550-590 nm, Em. = 608-683 nm), respectively. For confocal microscopy, fluorescence images were acquired using AX-Ti2 system.

### Microplate reader

Fluorescence intensities of samples loaded into 384-well plates (Greiner Bio-One, 784900) were measured using an EnVision microplate reader (PerkinElmer) with the FITC filter settings.

### Single-molecule enzyme activity assay

Digital enzyme assays were performed using commercially available microdevices^6,7^ (Simoa disk; Quanterix). A 35 µL of mixture of enzyme and reagents in buffer was loaded into the microdevice by manual pipetting. Then, 50 µL of FC-70 (TCI) was introduced into the device to flush out excess reaction mixture. After incubation at 25°C, the fluorescence images were acquired using an epifluorescence microscope (for LDH) and a confocal microscope (for PK).

### Image processing and data analysis

Images were processed using the GA3 module of NIS Elements software (Nikon). First, all fluorescence images were background-corrected using a rolling ball correction (diameter = 5 μm). Fluorescence-positive ROIs were then identified in the FITC channel using bright-spot detection (diameter = 3 μm, contrast threshold = 220–250). Irregular fluorescent spots derived from debris or air bubbles were excluded based on ROI size and shape after ROI dilation and removal of overlapping ROIs. Separately, all analyzable chambers were identified in the mCherry channel using spot detection (diameter = 3 μm). The mean occupancy (λ) was calculated as the fraction of FITC-positive chambers based on the low-occupancy approximation λ ≈ p of the Poisson distribution^8–10^, where p represents the number of FITC-positive chambers divided by the total number of analyzable chambers identified in the mCherry channel. The threshold separating the LDHA-major and LDHB-major fractions was determined by comparison with the fluorescence intensity distributions of recombinant LDHA and LDHB measured under the same conditions. Signals on either side of this threshold were classified as the LDHA-major and LDHB-major fractions, respectively.

### Shoulder analysis

To quantify the LDHA-associated shoulder in single-molecule fluorescence intensity distributions, each histogram was fitted with a single Gaussian function representing the major LDH population. The excess signal above the fitted distribution on the high-intensity side was defined as the shoulder fraction. The shoulder ratio was calculated as the integrated area of this right-sided excess relative to the total probability density and expressed as a percentage. The analysis script is provided in the Supplementary Information.

### Preparation of T cells

Ethical approval for isolating mouse CD8^+^ T cells was obtained from animal research committee at Kyoto University (Med Kyo 26529). Six-week-old female C57BL/6N mice were purchased from Japan SLC. Single cell suspension of mouse spleen and inguinal/axillary lymph nodes were prepared by mechanical mincing, followed by lysis of red blood cells using ACK Lysing Buffer (Thermo Fisher Scientific). Naïve CD8^+^ T cells were initially enriched using the MojoSort Mouse CD8 naïve e T Cell Isolation Kit (Biolegend) according to the manufacturer’s instructions. Cells were stained with anti-CD8 mAb-APC (clone: 53-6.7), anti-CD44 mAb-FITC (clone: IM7), and anti-CD62L mAb-APC (clone: MEL-14) (all from Biolegend) and sorted on BD FACSymphony S6 (BD Biosciences). Naïve CD8^+^ T cells were identified and sorted as CD8 positive, CD44 low, CD62L high (purity >98%). For stimulation, flat bottom 96-well plates (Thermo Fisher Scientific) were pre-coated with anti-CD3 mAb (Biolegend, clone: 145-2C11, 1 μg/mL) and anti-CD28 mAb (Biolegend, clone: 37.51, 1 μg/mL) at 37°C for 4 h. Isolated Naïve CD8^+^ T cells were seeded at 1×10^5^ cells/well in 250 μL RPMI culture media (supplemented with 10% FCS, non-essential amino acid solution and Penicillin/Streptomycin) with recombinant mouse IL-2 alone (Peprotech, 10 ng/mL) or in combination with mouse recombinant IL-12 (Peprotech, 1 ng/mL) for 48 h.

### Preparation of single-cells using FACS and proteoform analysis

Naïve CD8^+^ T cells (as indicated above) and stimulated CD8^+^ T cells (stained with anti-CD8 mAb-APC and Zombie NIR Fixable Viability Kit (Biolegend)) were sorted using FACSymphony S6 (BD Biosciences). Sorted cells were directly loaded into 96 well V bottom plate (Thermo Fisher Scientific) filled with 10 μL lysis buffer (HEPES-Na buffer (10 mM, pH 7.4) containing Triton X-100 (1.5 mM)). Lysate was flash frozen using liquid N_2_ and stored at 313:14280°C. For frozen lysate, 30 μL dilution buffer (HEPES-Na buffer (100 mM, pH 7.4) containing CaCl_2_ (1 mM), MgCl_2_ (1 mM) and DTT (100 μM)) warmed to 37°C and 10 μL solution of iLACCO (50 μM), pyruvate (10 mM), and NADH (5 mM) in dilution buffer was added and the mixture was loaded into microdevice. The microdevice was sealed using 50 μL FC-70 and fluorescence images were acquired after incubating at 25°C for 40 min.

### Cell culture

All cells were purchased from ATCC. A549 lung cancer cells and MDA-MB-231 breast cancer cells were cultured in RPMI containing 10% fetal bovine serum (FBS) and 1% penicillin-streptomycin (PS). Hep G2 liver cancer cells, MIA Paca-2 pancreatic cancer cells, HT-29 colorectal cancer cells and B16-F10 melanoma cells were cultured in Dulbecco’s modified Eagle medium (DMEM) containing 10% fetal bovine serum (FBS) and 1% penicillin-streptomycin (PS).

### Lysate preparation and measurement of extracellular release of LDH from culture cells

Cells cultured in 10-cm dishes were washed three times with phosphate buffered saline (PBS, Wako). Cells were then suspended in 1.5 mL PBS by scraping, transferred to 1.5-mL tubes, and collected by centrifugation at 800 × *g* for 1 min. The supernatant was discarded, and the cells were washed three times with Hank’s balanced salt solution (HBSS, 800 × *g*, 1 min) and resuspended in 300 μL HBSS. Cell counting was performed using a cell counter (Countess 3, Thermo Fisher Scientific). For preparation of cell lysates, 100 μL of the cell suspension was mixed with an equal volume of 2× lysis buffer (10 mM HEPES, pH 7.4, 3 mM Triton X-100) and incubated at 25°C for 10 min. The lysate was centrifuged at 12,000 × *g* for 1 min, and the supernatant was collected for subsequent analysis. To evaluate extracellular LDH release, cellular suspensions were incubated at 37°C, and extracellular fractions were collected at 0 and 1 h by centrifugation at 5,000 × *g* for 1 min.

### Statistics

Statistical analysis was not performed in this study. Data from replicate experiments are shown with error bars representing the standard deviation (S. D.), with the number of replicates (n) indicated in the figure legends. Values in tables are presented as mean ± S. D.; where values were derived from multiple measured quantities, the S. D. was calculated by error propagation.

## ASSOCIATED CONTENT

### Supplementary Information

Methods, supplementary figures and supplementary references.

### Data availability

Source data underlying all graphs in this study are provided with the paper as an Excel file. All other data supporting the findings of this study are available from the corresponding author upon request.

### Code availability

The custom Python code used for shoulder fraction analysis is provided in the Supplementary Information.

## AUTHOR INFORMATION

### Author Contributions

T.K. conceived and supervised the study. T.K. and R.E.C. designed the biosensor-based SEAP assay. S.I. prepared and characterized the fluorescent protein biosensors. T.K. and M.M. developed and performed the biosensor-based SEAP assay. M.T. and Y.S. designed the T-cell experiments and prepared the samples. T.K., T.I., R.T., R.K., H.K., and T.M. performed the animal experiments. Y.K. assembled and prepared the human plasma samples for the first cohort. K.H. assembled and prepared the human plasma samples for the second cohort. T.K. and T.M. developed the quantitative models and analyzed the data. T.T., M.T., Y.S., Y.K., K.H., R.K., Y.U., H.K., T.M., and R.E.C. contributed to data interpretation. T.K. wrote the manuscript with input from all authors.

## Acknowledgements

We thank Mr. Naoki Seike and Dr. Saaya Hario for technical support on protein sample preparation, Mr. Manato Kamiya, Ms. Kanon Sasao, and Ms. Yoshiko Nakajima for technical support in data acquisition, and Dr. Norimichi Nagano and Ms. Masami Kawana for technical support in plasma sample collection. This work was financially supported by JSPS (20H04694, 21A303, 22H02217, 23K23484, 25K01911 and 25K22520 to T. K., 24K10092 to M.T., 23K26794, 26H01674, and 26K01641 to T.T., and 24H00489 and 24H02267 to R.E.C.), JST (PRESTO (13414915), PRESTO Network (17949814) and FOREST (24012649) to T.K., START (20353017) to T.K., T.M., and K.H., CREST (JPMJCR25T3) to R.E.C., and ACT-X (JPMJAX2532) to S.I.), and AMED (FORCE (22581634) and P-PROMOTE (25131640) to T.K. T.M. and K.H., Research on Development of New Drugs (23809006) for T.K. and T.M., P-PROMOTE (18cm0106403h0003) and P-CREATE (25ama221431h0002) to K.H). Work in the lab of T. K. was supported by The Naito Foundation, The Mochida Memorial Foundation for Medical and Pharmaceutical Research, Chugai Foundation for Innovative Drug Discovery Science, MSD Life Science Foundation, Hoansha Foundation, and The University of Tokyo Gap Fund Program. Work in the lab of R.E.C. was supported by The Mitsubishi Foundation, The Precise Measurement Technology Promotion Foundation.

