## Supplementary_Information for "Biosensor-enabled single-molecule enzyme activity profiling for function-based molecular counting"

##### Contents

Supplementary tables and figures

Python codes

Supplementary references

##### Supplementary tables and figures

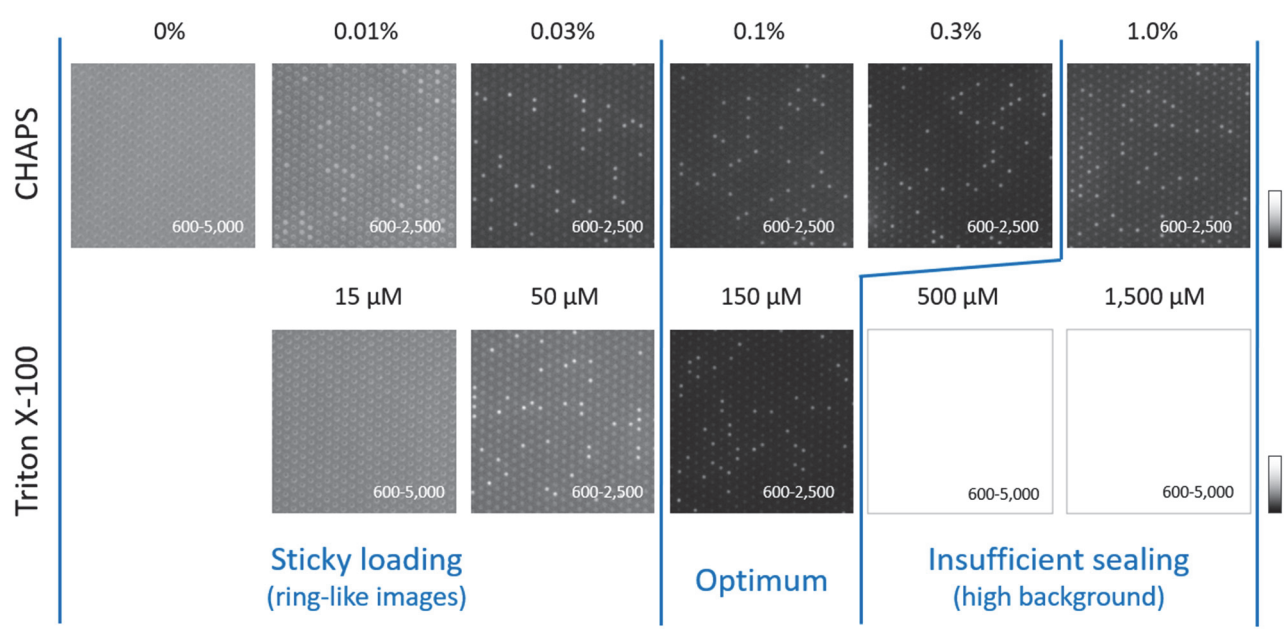

**Figure S1. Optimization of detergents for loading protein biosensors into microdevice.**

LDHB (0.1 ng/mL) was mixed with iLACCO (10 μM), pyruvate (1 mM) and NADH (1 mM) in HEPES-Na buffer (100 mM, pH 7.4) containing $\text{CaCl}_2$ (1 mM), $\text{MgCl}_2$ (1 mM), DTT (100 μM) and varied concentrations of detergents and incubated at 25°C for 45 min.

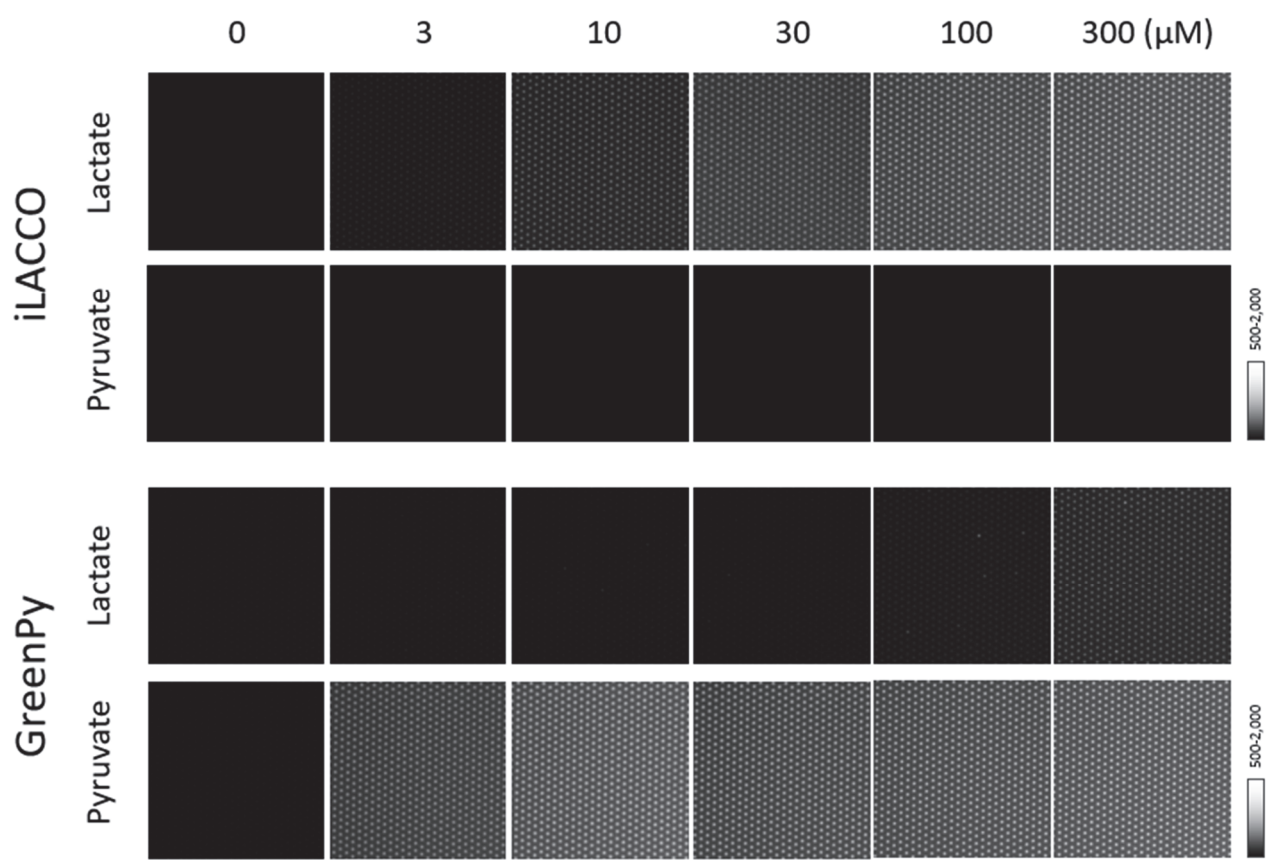

**Figure S2. Fluorescence responses of iLACCO and GreenPy in microdevice.**

iLACCO (10 $\mu\text{M}$) or GreenPy (10 $\mu\text{M}$) was mixed with varied concentrations of metabolites (L-lactic acid or pyruvic acid, 0-300 $\mu\text{M}$) in assay buffer (HEPES-Na buffer (100 mM, pH 7.4) containing $\text{CaCl}_2$ (1 mM), $\text{MgCl}_2$ (1 mM), DTT (100 $\mu\text{M}$), and Triton X-100 (150 $\mu\text{M}$)), and fluorescence images were acquired.

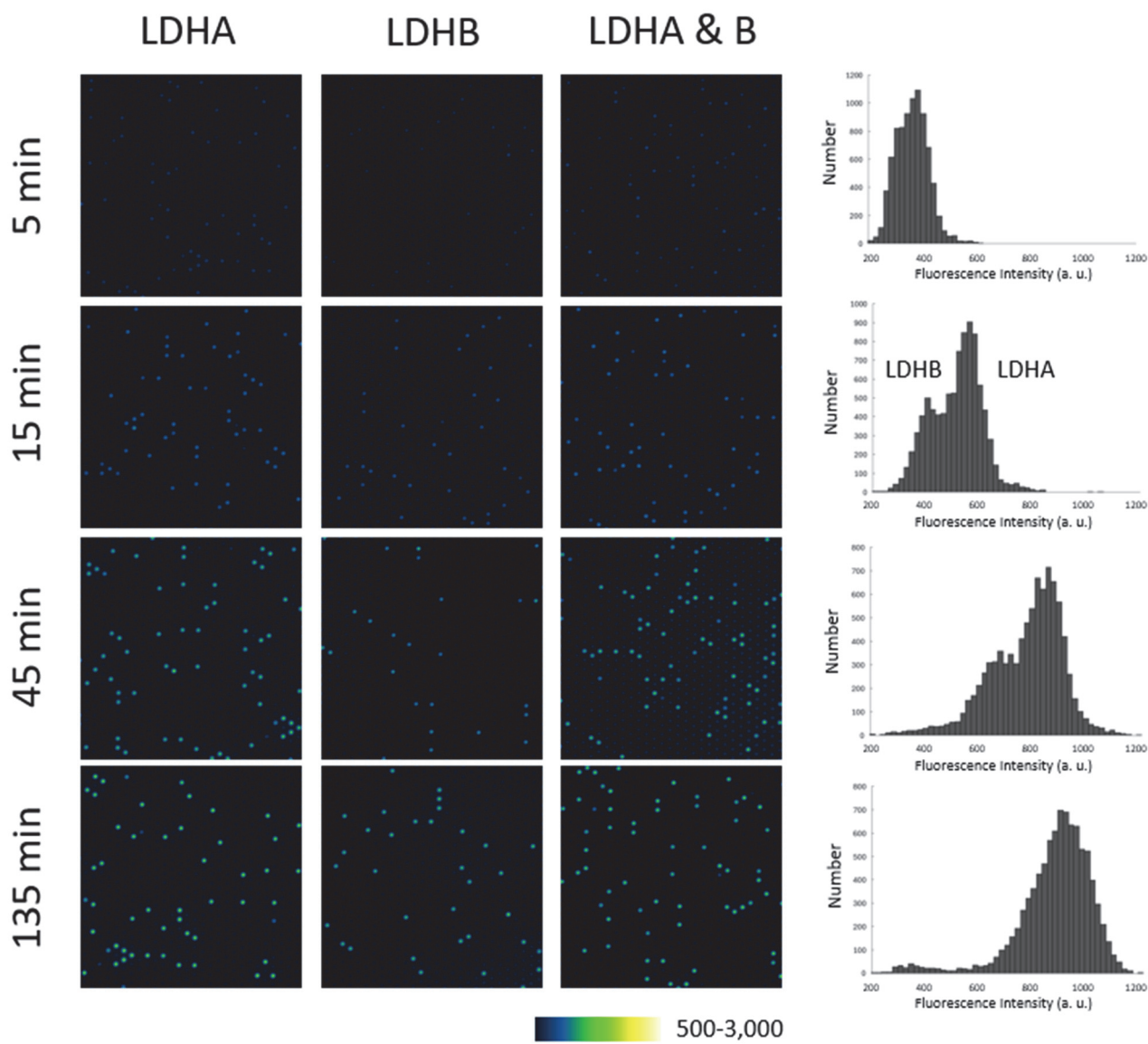

**Figure S3. Timecourse fluorescence changes of microchambers containing recombinant LDHA and LDHB.**

LDHA (1 ng/mL) and/or LDHB (0.1 ng/mL) were mixed with iLACCO (10 μM), pyruvate (2 mM) and NADH (1 mM) in assay buffer and incubated at 25°C for 5 to 135 min. The histogram showing the intensity distribution of bright spots corresponding to LDHA + B is shown.

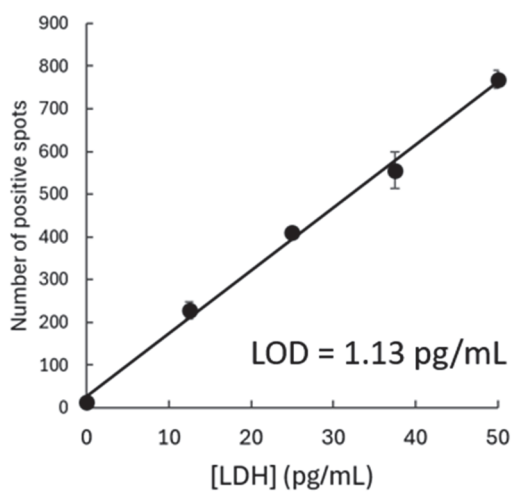

**Figure S4. Confirmation of the linearity of LDH spot number in microdevice-based assay.**

The measurement was performed by mixing LDHB (0-50 pg/mL), iLACCO (10 μM), pyruvate (2 mM) and NADH (1 mM) in assay buffer, loading into microdevice, and incubated at 25°C for 45 min. $n = 8$ for 0 pg/mL, and $n = 4$ for 12.5-50 pg/mL. Error bars represent S. D. The limit of detection (LOD) was calculated as the cell equivalent corresponding to the mean signal of the blank plus three standard deviations (3 S. D.).

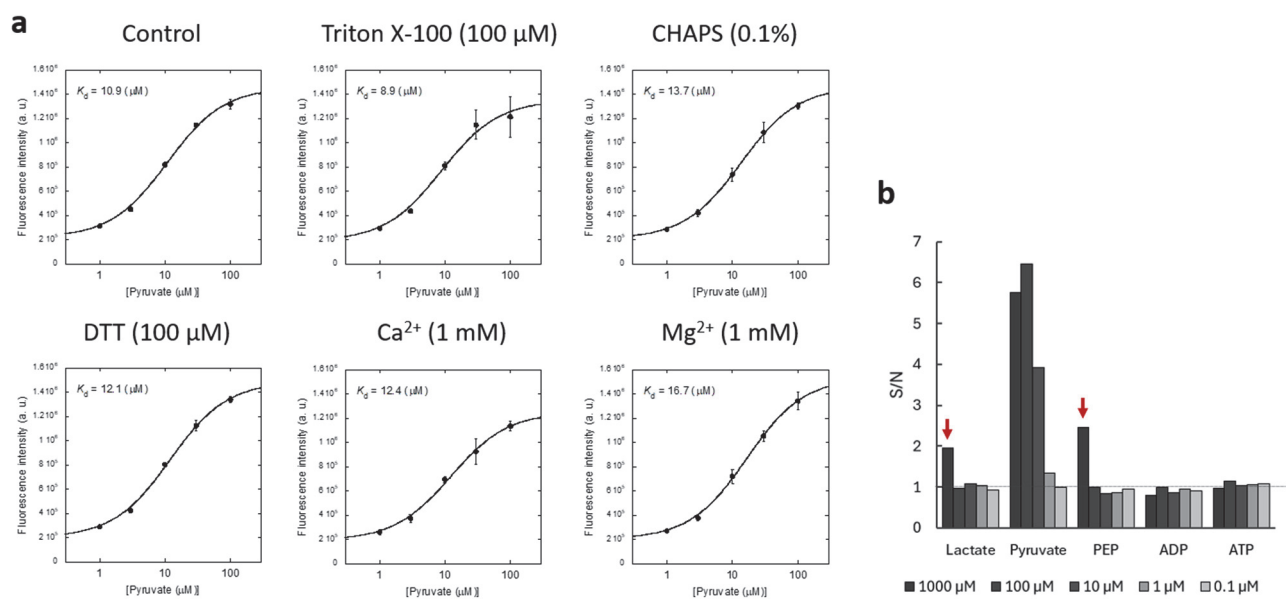

**Figure S5. Characterization of GreenPy under assay conditions.**

(a) Fluorescence response of GreenPy (10 μM) to pyruvate (1-100 μM) in HEPES-Na buffer (100 mM, pH 7.4, control) containing Triton X-100 (100 μM), CHAPS (0.1%), DTT (100 μM), $\text{CaCl}_2$ (1 mM), or $\text{MgCl}_2$ (1 mM). Apparent $K_d$ values determined from the fitted concentration-response curves are indicated in each panel. Data are shown as mean ± S. D. ($n = 3$). (b) Selectivity of GreenPy toward lactate, pyruvate, PEP, ADP, and ATP at the indicated concentrations (0.1-1000 μM). Fluorescence responses are expressed as signal-to-noise ratios (S/N) compared with the fluorescence intensity acquired without additional metabolites.

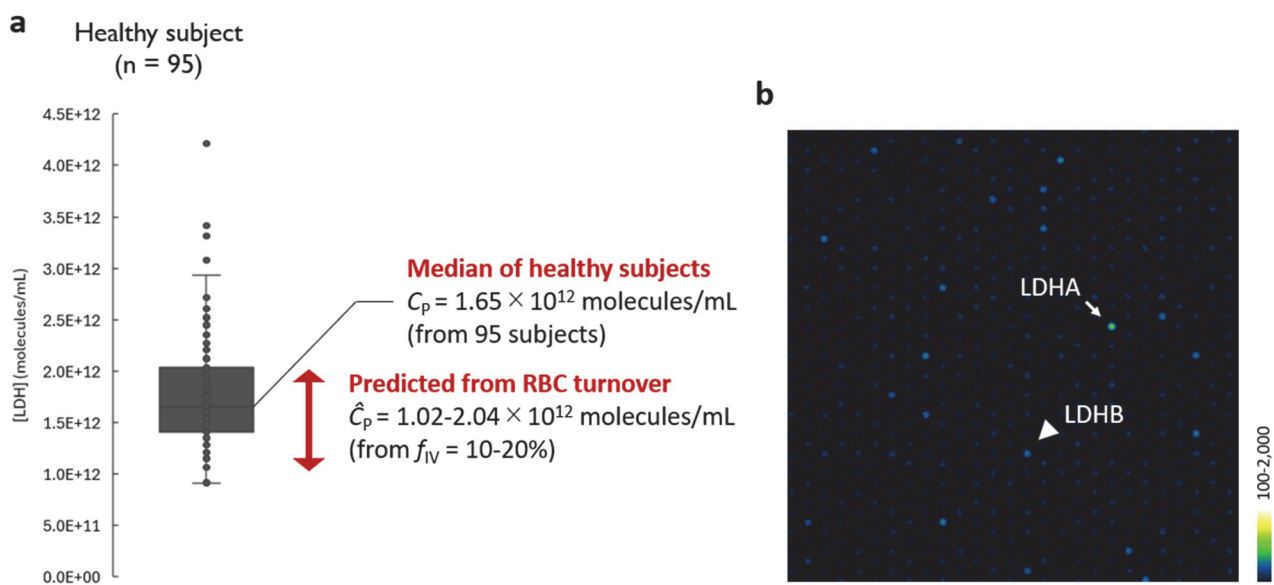

**Figure S6. Measurement of LDH activity of blood samples of healthy human subjects.**

(a) Distribution of amount of LDH number (molecules/mL) in 95 healthy human subjects. (b) Representative image in the analysis of plasma samples of healthy human subjects. The activity species of LDHA and LDHB were assigned from the comparison with the recombinant enzymes.

| Species |  |  | Human |  |  | Mouse |  |  | Rat |  |  |
| --- | --- | --- | --- | --- | --- | --- | --- | --- | --- | --- | --- |
| Details |  |  | 35-50 yo, male |  |  | C57BL/6J]cl, 6-7 wo, male |  |  | JCl:Wistar, 8 wo, male |  |  |
| Major species |  |  | LDHB |  |  | LDHA |  |  | LDHA |  |  |
| $\hat{C}_P = C_R * T_R * f_{IV} * \tau_{1/2} / (V_P * \ln 2)$ | | unit | Value | S. D. | Source | Value | S. D. | Source | Value | S. D. | Source |
| $C_R$ | RBC LDH expression | molecules/cell | 4.12E+04 | 6.34E+03 n = 4 | | 2.45E+04 | 3069 n = 4 | | 5.21E+04 | 6.66E+03 n = 3 | |
| $T_R$ | RBC Turnover | cells/d | 2.10E+11 | | [1] | 6.86E+08 | 6.79E+07 [2] | | 2.91E+09 | 2.56E+08 [2] | |
|  |  | cells/h | 8.75E+09 |  |  | 2.86E+07 | 2.83E+06 |  | 1.21E+08 | 1.06E+07 |  |
| $f_{IV}$ | Fraction of intravascular death | % | 13.3% | | [3] | 13.3% | | [3], [4] | 13.3% | | [3], [4] |
| $\tau_{1/2}$ | Circulating LDH half-life | h | 58.7 | 9 [5] | | 2 | | [6] | 8 | | [6] |
| $V_P$ | Plasma volume | mL | 3000 | | [3] | 1.41 | | | 12.71 | | |
| $C_P$ | Plasma LDH concentration | molecules/mL | 1.80E+12 | 5.96E+11 n = 95 | | 3.10E+11 | 6.53E+10 n = 5 | | 9.37E+11 | 2.48E+11 n = 5 | |
| $\hat{C}_P$ | Estimated plasma LDH concentration | molecules/mL | 1.35E+12 | 2.94E+11 | | 1.90E+11 | 3.04E+10 | | 7.64E+11 | 1.18E+11 | |
| $\hat{C}_P / C_P$ | | % | 75.2% | 29.8% | | 61.4% | 16.2% | | 81.5% | 25.0% | |
| [1] Sender et al. *Nat. Med.* **2021**<br>[2] Calculated as $T_R = C_{RBC} / \tau_{RBC}$<br>[3] Garby et al. *J. Clin. Invest.* **1959**<br>[4] Drvenica et al. *Biomolecules* **2022**<br>[5] Boyd et al. *Biochim. Biophys. Acta* **1967**<br>[6] Mahy et al. *Science* **1965**<br>The value of $f_{IV}$ (13.3%) was reported for humans (Garby et al. *J. Clin. Invest.* **1959**). The authors noted that, under physiological conditions, approximately 80–90% of erythrocytes are eliminated without releasing hemoglobin into the plasma through extravascular hemolysis. The experiment was performed in sheep. A study in humans (Bär et al., *Klin. Wochenschr.* **1970**) reported a half-life of $110 \pm 60$ h; however, this estimate was subsequently questioned by Smith et al. (*Clin. Chem.* **1987**), who suggested that the conventionally accepted half-life of approximately 110 h for serum LD-1 activity may substantially overestimate the actual LD-1 half-life in many patients after myocardial infarction. The half-life used here was calculated from $\alpha_2 = 0.0118 \pm 0.0018 \text{ h}^{-1}$. More than 60% of LDH5 disappeared within 3 h. | | | | | | | | | | | |
| $T_R = C_{RBC} / \tau_{RBC}$ | | unit | Value | SD | Source | Value | SD | Source | Value | SD | Source |
| $C_{RBC}$ | RBC concentration | cells/mL | | | | 1.09E+10 | 5.40E+08 [7] | | 8.70E+09 | | [8] |
| $W$ | Body weight | g | | | | 21.1 | 0.6 | | 324.3 | 8.5 | |
| $V_P$ | Plasma volume | mL/g body weight | | | | 0.067 | 0.004 [9] | | 0.0392 | 0.0032 [10] | |
|  | Plasma volume | mL |  |  |  | 1.4 |  |  | 12.7 |  |  |
| $V_B$ | Blood volume | mL/g body weight | | | | 0.121 | 0.008 [9] | | 0.0619 | 0.004 [10] | |
| $\tau_{RBC}$ | RBC half-life | d | | | | 40.7 | 1.9 [11] | | 60 | 3.2 [11] | |
| $T_R$ | RBC turnover | cells/d | | | | 6.86E+08 | 6.79E+07 | | 2.91E+09 | 2.56E+08 | |
| [7] Nemzek et al. *Inflamm. Res.* **2001**<br>[8] Stammers et al. *J. Physiol.* **1926**<br>[9] Kaliss et al. *Exp. Biol. Med.* **1950**<br>[10] Lee et al. *J. Nucl. Med.* **1985**<br>[11] Putten et al. *Blood* **1958** |  |  |  |  |  |  |  |  |  |  |  |

**Figure S7. Calculation of parameters $\hat{C}_P$ and $C_P$ using measured and referred values.**

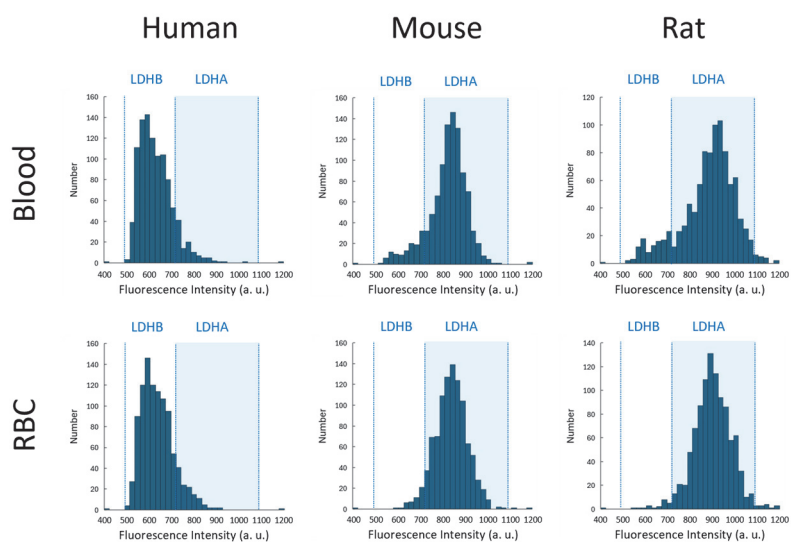

**Figure S8. Activity histograms of LDH species of plasma and erythrocyte (RBC) lysate of humans, mice, and rats.**

The measurement was performed by mixing plasma samples (1/2,500 dilution) or RBC lysate (1/10,000 dilution), iLACCO (10 μM), pyruvate (2 mM) and NADH (1 mM) in assay buffer, loading into microdevice, and incubated at 25°C for 45 min.

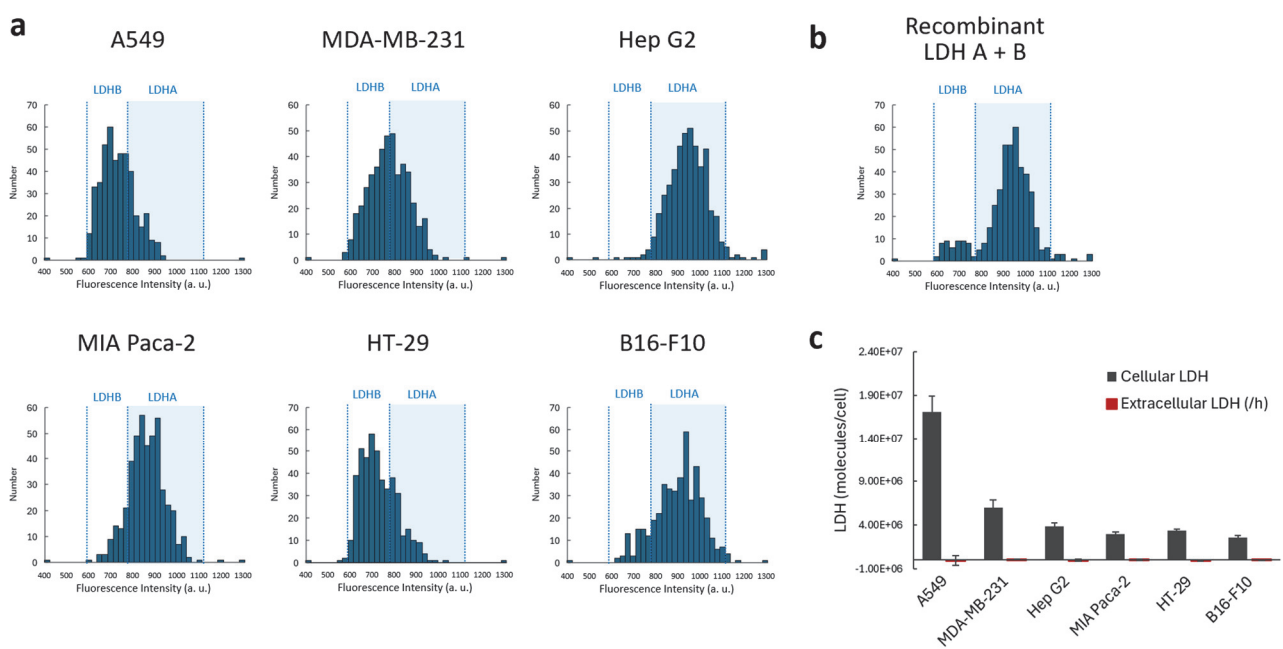

**Figure S9. Activity histograms of LDH species of cancer cell lysates.**

(a) Activity histograms of cancer cell lysate. The measurement was performed by mixing plasma samples (1/2,500 dilution) or cell lysate (1/10,000 dilution), iLACCO (10 μM), pyruvate (2 mM) and NADH (1 mM) in assay buffer, loading into microdevice, and incubated at 25°C for 45 min. (b) Activity histograms of mixture of LDHA (1 ng/mL) and LDHB (0.1 ng/mL) measured in the same condition as (a). (c) Quantification of cellular LDH and extracellular LDH release after incubating cells in Hank's balanced salt solution (HBSS) at 37°C for 1 h. The release was calculated as the differences between extracellular LDH at 1 h and at 0 h. Error bars represent S. D. (n = 3).

### Control

### Pancreatic cancer

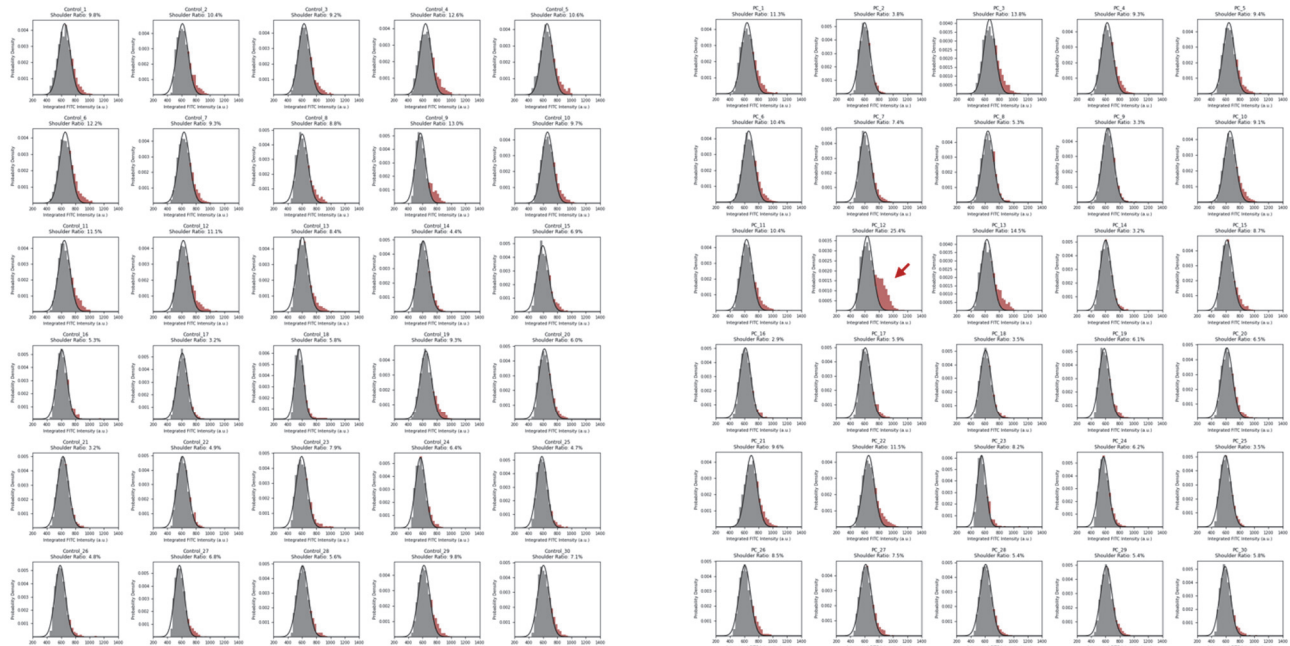

**Figure S10. Activity histograms of plasma from healthy subjects (control, n = 30) and patients with pancreatic cancer (n = 30).**

The measurement was performed by mixing plasma samples (1/2,500 dilution) with iLACCO (10 μM), pyruvate (2 mM) and NADH (1 mM) in assay buffer, loading into microdevice, and incubated at 25°C for 45 min. A red arrow indicates the patient having notable shoulder peak.

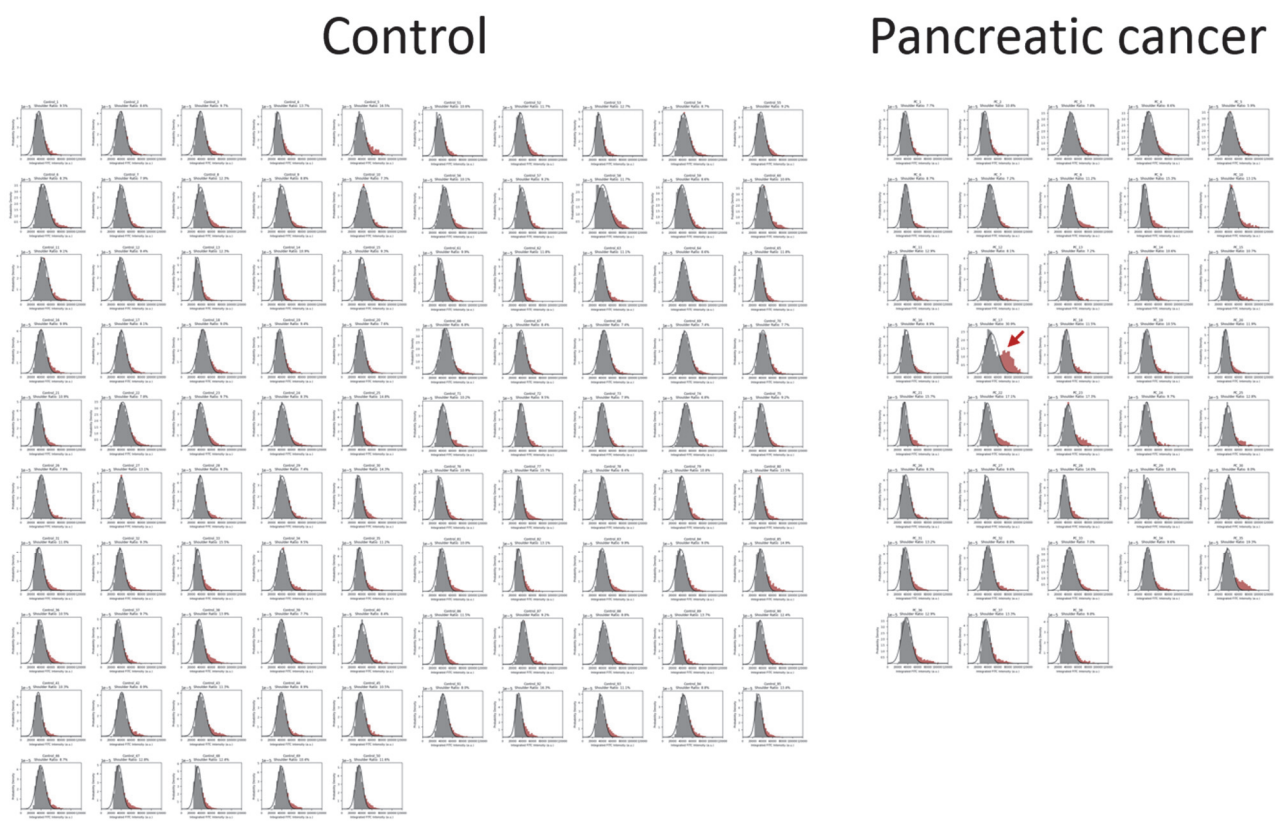

**Figure S11. Activity histograms of plasma from healthy subjects (control, n = 95) and patients with pancreatic cancer (n = 38) in 2<sup>nd</sup> cohort.**

The measurement was performed by mixing plasma samples (1/2,500 dilution) with iLACCO (10 μM), pyruvate (2 mM) and NADH (1 mM) in assay buffer, loading into microdevice, and incubated at 25°C for 45 min. A red arrow indicates the patient having notable shoulder peak.

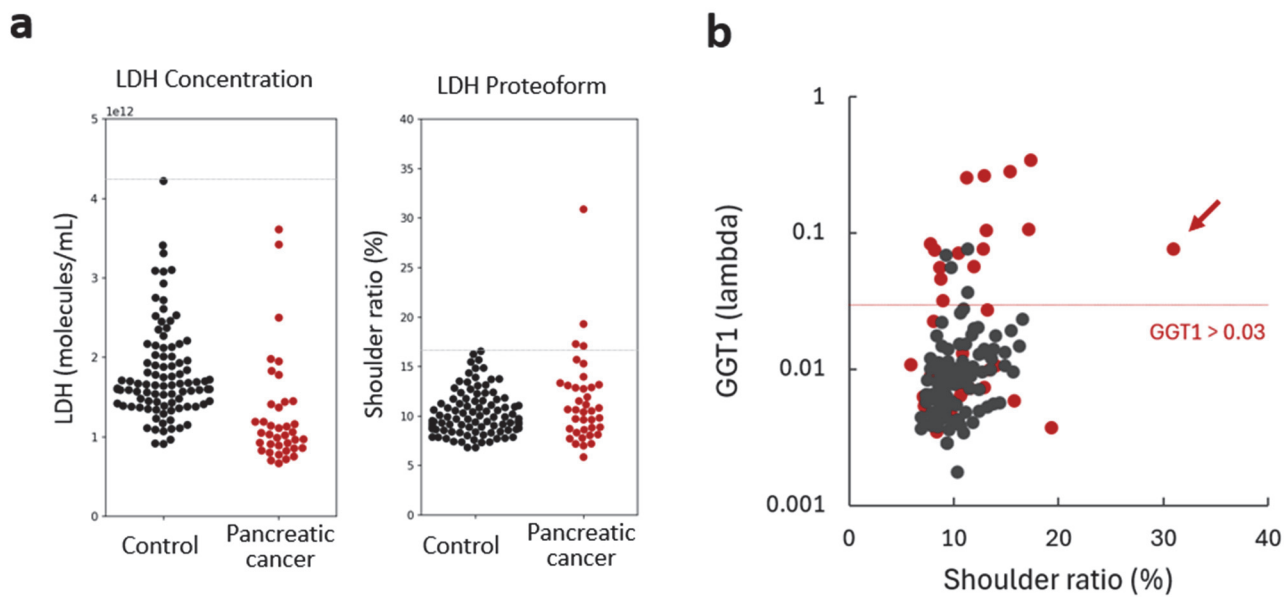

**Figure S12. Detailed analysis of data in 2<sup>nd</sup> cohort.**

(a) Results of total LDH number (left) or shoulder ratio (%) of 2<sup>nd</sup> cohort. $n = 95$ for healthy subjects and $n = 38$ for pancreatic cancer patients. (b) Correlation between shoulder ratio (%) and plasma GGT1 concentration<sup>11</sup>. The Red dots indicate patients with pancreatic cancer and the black dots indicate healthy subjects. A red arrow indicates the patient having notable shoulder peak. The red line indicates the GGT1 threshold ($> 0.03$) used to identify patients with pancreatic cancer accompanied by biliary obstruction<sup>11</sup>.

### Python code

#### Shoulder fraction analysis

The Python code used to quantify the shoulder fraction in single-molecule fluorescence intensity distributions is provided below.

```
import numpy as np
import pandas as pd
import matplotlib.pyplot as plt
from scipy.optimize import curve_fit
from google.colab import files

def gaussian(x, amplitude, mean, std_dev):
    return amplitude * np.exp(-((x - mean) ** 2) / (2 * (std_dev ** 2)))

# --- USER CONFIGURATION ---
MIN_VALUE = ...
MAX_VALUE = ...
BINS_NUM = ...
SIGMA_INIT = ...
SIGMA_MIN = ...
SIGMA_MAX = ...

# --- DATA INGESTION ---
print("Please upload your target CSV data file:")
uploaded = files.upload()
filename = list(uploaded.keys())[0]

try:
    df = pd.read_csv(filename, header=[0, 1])
    if "Unnamed" in str(df.columns[0][1]):
        df = pd.read_csv(filename, header=0)
except Exception:
    df = pd.read_csv(filename, header=0)

# --- BATCH ANALYSIS & FITTING ---
results = []
num_samples = len(df.columns)
n_cols = 5
n_rows = int(np.ceil(num_samples / n_cols))

fig, axes = plt.subplots(n_rows, n_cols, figsize=(15, 2.7 * n_rows))
axes = np.array(axes).reshape(-1)

for i, col in enumerate(df.columns):
    if isinstance(col, tuple):
        group_label, sample_id = str(col[0]), str(col[1])
        display_title = f"{group_label}_{sample_id}"
    else:
        display_title = str(col)

    raw_data = df[col].dropna()
    valid_data = raw_data[(raw_data >= MIN_VALUE) & (raw_data <=
MAX_VALUE)]
    total_count = len(valid_data)

    if total_count == 0:
        continue

    density, bin_edges = np.histogram(
        valid_data, bins=BINS_NUM, range=(MIN_VALUE, MAX_VALUE),
density=True)
    bin_centers = (bin_edges[:-1] + bin_edges[1:]) / 2
    bin_width = bin_edges[1] - bin_edges[0]

    peak_idx = np.argmax(density)
    initial_mu = bin_centers[peak_idx]
    initial_a = density[peak_idx]
    p0 = [initial_a, initial_mu, SIGMA_INIT]

    bounds = (
        [0, MIN_VALUE, SIGMA_MIN],
        [np.inf, MAX_VALUE, SIGMA_MAX]
    )

    try:
        popt, _ = curve_fit(
            gaussian, bin_centers, density, p0=p0, bounds=bounds,
maxfev=5000
        )
        a_fit, mu_fit, sigma_fit = popt
        fit_success = True
    except Exception:
        fit_success = False
        mu_fit, sigma_fit = np.nan, np.nan

    if fit_success:
        fitted_curve = gaussian(bin_centers, *popt)
        excess_density = np.maximum(0, density - fitted_curve)
        right_side_mask = bin_centers >= mu_fit
        shoulder_density = np.where(right_side_mask, excess_density, 0)

        shoulder_fraction = np.sum(shoulder_density * bin_width)
        shoulder_percentage = shoulder_fraction * 100
        shoulder_count = int(np.round(total_count * shoulder_fraction))
    else:
        shoulder_percentage = np.nan
        shoulder_count = np.nan

    results.append({
        "Sample_ID": display_title,
        "Total_Particles": total_count,
        "Peak_Center_mu": round(mu_fit, 2) if fit_success else np.nan,
        "Peak_Width_sigma": round(sigma_fit, 2) if fit_success else np.nan,
        "Shoulder_Particles": shoulder_count if fit_success else np.nan,
        "Shoulder_Ratio_pct": round(shoulder_percentage, 2) if fit_success
    else np.nan
    })

    ax = axes[i]
    ax.bar(
        bin_centers, density, width=bin_width,
        color='dimgray', alpha=0.75, edgecolor='none'
    )

    if fit_success:
        ax.bar(
            bin_centers, shoulder_density, width=bin_width,
bottom=fitted_curve,
            color='indianred', alpha=0.75, edgecolor='none'
        )
        x_smooth = np.linspace(MIN_VALUE, MAX_VALUE, 300)
        y_smooth = gaussian(x_smooth, *popt)
        ax.plot(x_smooth, y_smooth, color='black', linewidth=1.2)
        ax.set_title(f"{display_title} Shoulder Ratio:
{shoulder_percentage:.1f}%", fontsize=8.5)
    else:
        ax.set_title(f"{display_title} [Fit Failed]", fontsize=8.5)

    ax.set_xlim(MIN_VALUE, MAX_VALUE)
    ax.set_xlabel("Fluorescence Intensity (a.u.)", fontsize=7.5)
    ax.set_ylabel("Probability Density", fontsize=7.5)
    ax.tick_params(labelsize=7)
    ax.grid(False)

for j in range(i + 1, len(axes)):
    fig.delaxes(axes[j])

plt.tight_layout()
plt.show()

# --- OUTPUT EXPORT ---
results_df = pd.DataFrame(results)
display(results_df)

output_csv_path = "summary_shoulder_analysis.csv"
results_df.to_csv(output_csv_path, index=False, encoding='utf-8-sig')
files.download(output_csv_path)
```

#### Parameters used for analysis

For the first cohort, MIN\_VALUE = 400, MAX\_VALUE = 1200, BINS\_NUM = 40, SIGMA\_INIT = 60, SIGMA\_MIN = 5, and SIGMA\_MAX = 80. For the second cohort, MIN\_VALUE = 10000, MAX\_VALUE = 120000, BINS\_NUM = 40, SIGMA\_INIT = 2500, SIGMA\_MIN = 100, and SIGMA\_MAX = 12500.
